# Androgen receptor alpha regulates aromatase expression in the ventromedial hypothalamus of male cichlids

**DOI:** 10.1101/2023.08.25.554697

**Authors:** Mariana S. Lopez, Beau A. Alward

**Affiliations:** University of Houston, Department of Psychology; University of Houston, Department of Biology and Biochemistry

## Abstract

Social behaviors are regulated by sex steroid hormones such as androgens and estrogens. However, the specific molecular and neural processes modulated by steroid hormones to generate social behaviors remain to be elucidated. We investigated whether some actions of androgen signaling in the control of social behavior may occur through the regulation of estradiol synthesis in the highly social cichlid fish *Astatotilapia burtoni*. Specifically, we examined the expression of *cyp19a1*, a brain-specific aromatase, in the brains of male *A. burtoni* lacking a functional ARα gene, which was recently found to be necessary for aggression in this species. We found that *cyp19a1* expression is higher in wild-type males compared to ARα mutant males in the homolog of the mammalian ventromedial hypothalamus (VMH), a brain region that governs aggression across taxa. Using *in situ* hybridization chain reaction (HCR), we determined that ARα^+^ and *cyp19a1*^*+*^ cells are commonly nearby, but very infrequently was co-expression observed, including in the fish VMH. We speculate that ARα may modulate *cyp19a1* expression in the fish VMH to govern aggression through an indirect, potentially transsynaptic, mechanism. These studies provide novel insights into the hormonal mechanisms of social behavior and lay a foundation for future functional studies.

## Introduction

Steroid hormones regulate physiology and behavior via ligand-dependent receptor activation ^1,2^. Specifically, steroid hormones exert their effects via specific steroid receptors that when bound, will dimerize, and then translocate to the nucleus to alter the expression of suites of genes, thus altering cellular processes that give rise to modifications in biological functions. Additionally, steroid receptors can modulate cellular functions differently from modulating gene expression, by rapidly affecting neural processes via ion channels and secondary messenger cascades^3^. Despite the known link between steroid hormones, steroid receptors, cellular processes, and behavior, it remains unclear which genes in the brain steroids modify to alter behavior. This is in stark contrast to our knowledge of other steroid-sensitive tissues like the prostate, where some of the most comprehensive data on steroid receptor controlled gene expression are available^4–6^. A comprehensive understanding of the functions of steroid hormones in the regulation of the brain and behavior requires approaches that can elucidate the downstream genes controlled by steroid receptors.

Steroid receptors are expressed widely in the brain, including in areas of the brain known to control social behavior. Indeed, sex steroid receptors, including androgen receptors (AR) and estrogen receptors (ER), are expressed in regions of the social behavior network (SBN), a highly conserved, interconnected set of brain regions in vertebrates that respond to social cues and coordinate adaptive physiological and behavioral responses ^7^. These regions include the preoptic area (POA), the anterior hypothalamus (AH), the VMH (ventromedial hypothalamus), the lateral septum (LS) and BNST (bed nucleus of the stria terminalis)^8^. Given its conserved functions, the role of the SBN and functions of steroid hormone actions throughout have been extensively studied across diverse taxa to reveal fundamental insights into the neuroendocrine regulation of behavior.

One such species in which important insights have been gained on the hormonal regulation of behavior and SBN functions is the African cichlid fish *Astatotilapia burtoni. A. burtoni* is a genetically tractable and highly social fish species, making it an ideal organism to study the hormonal and neural mechanisms of social behavior^9^. Male *A. burtoni* form a flexible social hierarchy in which certain males are dominant (DOM) and others are subordinate (SUB). DOM males are brightly colored, have large testes, produce territorial and reproductive behaviors, and have increased levels of sex steroid hormones including testosterone, 11-ketotestosterone, and estradiol ^10^.SUB males are drably colored, produce subordinate behaviors like fleeing, are less likely to reproduce, have smaller testes, and have lower levels of sex steroid hormones. Importantly, males can switch rapidly between these two social states depending on the social context ^9,11,12^. Female *A. burtoni* also perform territorial and reproductive behaviors but are not known to naturally form a social hierarchy.

Previous research in male *A. burtoni* shows that gene expression in the brains of DOM and SUB males also differs. In fact, DOM males have higher levels of immediate early gene and sex steroid receptor expression in the SBN compared to ND males. Additionally, male fish that are ascending, or rising in rank from SUB to DOM social status, show increased expression of these genes in the SBN^10^. Taken together, this indicates that hormones and the mechanisms associated with behavioral production and physiological regulation in response to the environment, are all tightly linked and can change quickly depending on environment and social cues.

Recently it was discovered that androgen receptors (ARs) control distinct traits of social dominance in male *A. burtoni*. CRISPR/Cas9 gene editing was used to engineer mutant *A. burtoni* that lack one allele (heterozygous or Het mutants) or both alleles (homozygous null or knockout (KO) mutants)) encoding ARα or ARβ. Male AR mutants were then tested for traits associated with social dominance including aggressive and courtship displays, bright coloration, and testes mass^13,14^. Both ARα Het and KO mutants performed fewer aggressive, and courtship displays compared to wild type (WT) males and were no different from one another on these measures. Coloration and testes mass within ARα mutants, however, was not significantly reduced. On the other hand, ARβ heterozygous and KO mutants lacked bright coloration and possessed reduced testes mass compared to WT males but produced normal levels of aggressive and courtship behavior. Therefore, ARs in *A. burtoni* regulate distinct aspects of social dominance.

Previous research indicates the conversion of androgens to estrogens by the enzyme aromatase is crucial for the activation of aggressive displays in male *A. burtoni*. O’Connell and Hofmann^15^ injected male *A. burtoni* with 17β-estradiol and observed an increase in aggression, while injection of an estrogen receptor (ER) antagonist caused a reduction in aggression. In subsequent work, male *A. burtoni* were injected with the aromatase inhibitor fadrozole. This manipulation caused a significant decrease in aggression compared to males injected with just vehicle^16^. Hence, in male *A. burtoni* androgenic signaling via ARα, as well as the synthesis and action of estradiol, may be required for performing aggressive behavior. Furthermore, in contrast to other vertebrate species, in which only one aromatase gene is present, there are two different copies of the aromatase gene in teleost species, one primarily expressed in the gonads called *cyp19a1a* and the other expressed primarily in the brain called *cyp19a1*^17–19^.

Research in other species has shown that androgenic signaling enhances aromatase expression in the brain. Castrated male mice have lower aromatase expression in the diencephalon compared to intact male mice, while castrated male mice treated with testosterone— but not estradiol—show an increase in aromatase expression in the diencephalon to intact levels^20^. The androgen dehydroepiandrosterone (DHEA) increases aromatase expression in the preoptic area (POA) and ventromedial hypothalamus (VMH) of male song sparrows (*Melospiza melodia)*^21^. In male rats, aromatase expression in the POA and mediobasal hypothalamus, which contains the VMH, was lower in castrated males compared to intact males. However, treatment with testosterone or dihydrotestosterone (DHT), but not estradiol, in castrated males restored aromatase expression to intact levels^22^. Thus, the modulation of aromatase expression by androgens may be direct since aromatase-positive cell bodies in the hypothalamus co-expressed with AR in mice^23^.

Cross-species and genetic approaches to compliment the above work on the role of steroid hormones in controlling aromatase expression is necessary to achieve a comprehensive understanding of these mechanisms. In the current study, we tested whether ARα mutant *A. burtoni* possessed reduced *cyp19a1* expression in the brain. If this were the case, it would support the idea that ARα controls aggression via the upregulation of aromatase. Additionally, we used *in situ* hybridization chain reaction (HCR) to determine whether either AR and *cyp19a1* are expressed in the same cells in the brain. We hypothesized that 1) since ARα Het and KO performed significantly fewer aggressive displays compared to WTs and were no different than one another, they would both have lower *cyp19a1* expression compared to WTs in regions of the SBN; and 2) that ARα and *cyp19a1* would be expressed in the same cells in brain regions in the SBN.

## Materials and Methods

### Animal subjects and housing

The *A. burtoni* males used in this study for detection of *cyp19a1* expression were involved in another experiment in which behavior, body coloration, and gonad mass results have already been published, but for which brain gene expression has not been assessed^14^. Fish originated from a stock from Lake Tanganyika, Africa and were kept in laboratory conditions simulating their natural habitat (25°C; 12h day: 12h night cycle)^12,24^ in accordance with Association for Assessment and Accreditation of Laboratory Animal Care standards. All experimental procedures were approved by the Stanford University Administrative Panel for Laboratory Animal Care (Protocol #9882) and by the University of Houston Institutional Animal Care and Use Committee (Protocol #202000001).

#### ARα mutant males

We used male *A. burtoni* with a mutant ARα (accession number, NW_005179415) allele generated using CRISPR/Cas9 gene editing. The process used for generating these mutants is described in full detail in Alward et al^14^. The Cas9 enzyme was directed towards a site within exon 1 upstream from the DNA binding and ligand binding domains. Specifically, Cas9 was directed towards two regions within exon 1 using two single guide RNAs (sgRNAs) targeting sequence ARα-A, 5′-ACTGTGGCGGATACTTCTCG-3′ and sequence ARα-B, 5′-GGTGCGCAAACT-GTGACGCG-3′, whose cut sites were separated by 178 bp. G1 offspring from injected fish carried a frameshift deletion within exon 1 of ARα of 50-bp. We outcrossed this allele to unrelated wild-type fish and then intercrossed heterozygous mutants to obtain biallelic ARα mutants (ARα^d50/d50^, hereon referred to as ARα^KO^), Het ARα mutants (ARα^d50/+^, hereon referred to as ARα^Het^), and ARα wild types (ARα^+/+^, hereon referred to as ARα^WT^).

#### Housing: ARα mutants

Male offspring from AR^Het^ x AR^Het^ matings were housed in stable dominant tanks, wherein five to eight size-matched males are housed in a 121-liter tank with 10 to 15 females and five potential mating sites that are represented by halved terra cotta pots^14^. Housing the males used in this study in stable dominant tanks ensures 1) all fish came from a similar social environment and 2) were housed in an environment that reliably induces traits of social dominance, including elevated androgen levels^25–28^. In the previously published work on the fish used here, there were no differences in androgen levels between ARα^WT^, ARα^Het^, and ARα^KO^ males, and these values were well within the range typical of DOM males^14,25^. Males of each genotype were housed randomly and blind to the experimenter, as PCR-based genotyping was not performed until the harvesting of tissue at the end of the experiment.

Males were housed in stable dominant tanks for 4-12 weeks before being moved to a dyad assay set up to test DOM-typical behaviors that were assessed in the previously published work^14^. Upon removal from the stable dominant tank, fish underwent a pre-assay procedure that took approximately 45 seconds. Fish were photographed, a small (1-2 mm) piece of their caudal fin was clipped for subsequent DNA isolation and genotyping, and then they were immediately moved to a 30-liter tank where they were housed with 3 smaller females and 1 smaller stimulus male. A camcorder (Canon VIXIA HF R80) mounted on a tripod was placed in front of their tank and recordings were made on the day they were added to the tank, then for two subsequent days. At 2 pm on the day after the second day, focal fish were removed, and tissue was harvested for physiological measurements and blood was collected. Brains from 15 males were used for the current experiment (ARα^WT^ n=5, ARα^HET^ n= 5, ARα^KO^ n= 5).

#### Housing: WT fish used for HCR

WT fish were housed for a 4–12-week period in an 80-gallon mixed-sex tank allowing for naturalistic social interactions to occur. Males were given the opportunity to rise in dominance by providing 3-5 potential mating sites and free access to females within the tank. Once the males became dominant as measured by enhanced reproductive activity, territoriality, and brightness in coloration, they were taken from their tank, dissected, and had tissue and blood harvested for physiological measurements. Similarly, female WT fish from the same community tank were allowed equal access to natural social interactions and access to males. Females were selected for dissection when they were in a gravid reproductive state as measured by observation of a distended abdomen assumed to be filled with eggs. Once the females met these criteria, they too were dissected, their brain and blood collected, and their physiological measures taken. Brains from two males and two females were collected and used for HCR.

### Tissue collection

Upon removal from their dyad assay tank, males from the ARα^Het^ cross were euthanized using rapid cervical transection, their brains and blood collected, and a fin clip taken for genotyping purposes. Fish used for HCR were removed from their home community tank and euthanized in an ice bath slurry before cervical transection, then their brains and blood collected. We measured standard length (SL) in cm from the tip of the mouth to the base of the tail fin. Body mass (BM) in grams was measured and recorded, and gonads were extracted and weighed. Blood for each animal was collected from the dorsal artery using 2-3 heparin coated capillary tubes (VWR^®^).

Brains for fish from the ARα^Het^ cross were dissected and fixed in 4% paraformaldehyde (PFA) with a pH of 7.4 for two hours before being cryoprotected using a sucrose gradient starting with 15% sucrose (in 1xPBS; Gibco) overnight and subsequently 30% sucrose (in 1xPBS; Gibco) for 1-2 days (or until sunk). Brains from WT fish used for HCR were fixed in 4% paraformaldehyde (PFA) with a pH of 7.0 for 24 hours before being cryoprotected using a sucrose gradient starting with 15% sucrose (in 1xPBS; Gibco) overnight and subsequently 30% sucrose (in 1xPBS; Gibco) for 1-2 days (or until sunk). Once all brains had sunk, they were embedded in Neg-50 (Epredia) and stored at -80°C until sectioning at 30μm on a cryostat (Thermo Scientific, HM525NX).

### Histology

#### In situ hybridization

*In situ* hybridization (ISH) of ARα mutant brains was performed to analyze the expression of the brain aromatase (*cyp19a1*) as previously described^29^. Briefly, PCR was used to amplify a sequence from *cyp19a1* using primers: *cyp19a1* forward, 5’-AGATGATAATCGCAGCCCCC-3’; *cyp19a1* reverse, 5’-TAATACGACTCACTATAGGGGAGTGACCAGGATGGCCTT -3’. The PCR product was subsequently confirmed using sanger sequencing and transcribed *in vitro* using T7 RNA polymerase (Promega) and Digoxygenin (DIG) labeled dNTP’s (Roche), the resultant RNA *cyp19a1* probe was then used for ISH.

Tissue for ISH was cryosectioned coronally at 30 μm (Thermo Scientific, HM525NX) at -20°C and mounted on Superfrost® slides (VWR) and allowed to dry for two hours before storage in - 80°C. To increase tissue adhesion, we coated slides with mounted tissue in a gelatin solution before starting the ISH. Specifically, gelatin (1mg/ml) was applied to each slide and placed on a slide warmer for 45 minutes at 50°C until the liquid evaporated. Subsequently the slides were dried in a desiccator overnight before continuing with ISH.

All steps of the ISH protocol were performed in a 40 ml coplin jar unless otherwise noted. ISH began by fixing the tissue in 4% PFA before adding Proteinase K (10mg/ml; Life technologies). After a second round of fixation with 4% PFA, sections were incubated in prehybridization buffer (50% formamide, 5X SSC, 0.1% Tween-20, 0.1% CHAPS, 5mM EDTA) for 2 hours at 62°C and subsequently incubated overnight at the same temperature in hybridization buffer (50% formamide, 5X SSC, 0.1% Tween-20, 0.1% CHAPS, 5mM EDTA, 1mg/ml torula yeast RNA, 100ug/ml Heparin, 1x Denhardt’s solution) with 32 ng of DIG-labeled RNA probe. Next day, slides were washed with 50% formamide (Thermo Fisher) and 2X SSC (Gibco) at 62°C. This was followed by three washes with 2X SSC at 37°C and treatment with RNaseA (200ng/ml; Qiagen) in 2X SSC at 37°C. The slides were then washed in Maleic acid Buffer (MABT; 100mM Maleic Acid, 150mM NaCl, 0.1% Tween-20), and subsequently blocked with MABT plus 2% BSA for 1.5 hours. Anti-DIG antibody fragments conjugated with alkaline phosphatase (Roche; 1: 5000) were diluted in MABT: 2%BSA and then incubated at 4°C overnight. On day 3, slides were washed with MABT and subsequently with alkaline phosphatase buffer (100mM Tris pH9.5, 50mM MgCl_2_, 100mM NaCl, 0.1% Tween-20, 5mM Levamisole (Tetramisole)) with NBT (37.5 mg /ml; Roche) and BCIP (94 mg /ml; Roche) for 12-16 hours at room temperature. Slides were then washed in 1x PBS, and cover slipped with Aquapolymount aqueous mounting media.

#### Hybridized Chain Reaction (HCR)

Tissue for HCR was cryosectioned at 30 μm at -20°C taking coronal sections that were mounted on Superfrost® slides (VWR). The mounted slides were allowed to dry for 24hrs at room temperature before being stored at -80°C before HCR. HCR was performed according to manufacturer protocols, recommendations, and reagents, unless otherwise noted^30^. In summary, tissue slides were exposed to LED light for 60mins to reduce autofluorescence. Subsequently, tissue was immersed in 0.2% Triton X-100 (in 1x PBS) (Thermo Fisher; Gibco) for 30 minutes to increase signal detection. Next, tissue was dehydrated with serial concentrations of EtOH (Thermo Fisher) at 50%, 70%, and 100%, before adding Proteinase K (1:2000) (Thermo Fisher) in 1xPBS. HCR probes were then hybridized to tissue by adding 16nMs of RNA probe in hybridization buffer for 12-16hrs at 37°C. The HCR probes used against ARα and ARβ were engineered (Molecular Instruments) using identical binding sequences from previous published work in *A. burtoni* ^31^. The HCR probe used against *cyp19a1* was made custom and is propriety information from the manufacturer (Molecular Instruments) and had zero complimentary binding to the paralogous gene *cyp19a1a*. Following the hybridization step, the tissue plus probe was exposed to probe amplifiers containing different fluorescent fluorophores per gene of interest, by adding 60nMs of fluorescent amplifier in amplification solution, and incubating at room temperature in a dark chamber overnight. Slides were then washed in 5X SSCT (0.1%Tween-20) (Gibco; Thermo Fisher), and cover slipped using Prolong Gold (Invitrogen) mounting media with DAPI. The fluorophores for *cyp19a1* and each of the ARs had excitation wavelengths in the GFP and CY5 range, respectively; thus, two series from the same brain were run in tandem allowing for the visualization of both *cyp19a1* with ARα and *cyp19a1* with ARβ. Due to the low signal intensity of both ARα and ARβ, we did not use autofluorescence quenching methods, which reduced signal intensity in optimization and validation of our HCR protocol. While this preserved signal intensity for reliable anatomical localization of ARα and ARβ signal, artifactual autofluorescence from what are believed to be blood vessels remained in many of the sections. They were verified to be artifacts by their presence in all wavelength filters and thus disregarded in all HCR runs.

### Microscopy and image analysis

#### Microscopy: Cyp19a1 in AR mutants using ISH

All images of stained *cyp19a1* ISH brains were obtained using Nikon eclipse 80i microscope and MicroFire ™ camera set. Images were taken at 4x, 10x, and 20x magnification, but only images at 4x were used for the quantification of *cyp19a1* signal. Consistent exposure settings were used for taking pictures across each of the magnifications as well as across different regions of the brain.

#### Microscopy: Co-expression of cyp19a1 and ARα and ARβ using HCR

All images of brains in which *cyp19a1*, ARα, and ARβ expression was detected using HCR were obtained using Nikon eclipse Ti2 microscope and DS-Qi2, Fi3 camera set. Images were taken at 10x and 20x magnification using the stitching function to get a complete picture of the coronal section of interest. Additionally, Z-stacks were taken at 30x magnification in the specific regions of interest allowing for visualization of possible co-expression of ARα and *cyp19a1* and ARβ and *cyp19a1*. Furthermore, the same exposure settings were consistently used for all the pictures taken to ensure consistency across all brain regions analyzed. Images depicting *cyp19a1* and ARβ were color changed using NIS elements (Nikon) software so that ARβ appeared magenta to distinguish from images depicting ARα and *cyp19a1*.

#### Quantification of cyp19a1 expression

We quantified *cyp19a1* expression in areas of the SBN. In the forebrain, we quantified expression of *cyp19a1* in the following regions of the SBN: the POA, subcomissural nucleus of the ventral telencephalon (Vs) a partial homolog of the BNST, and ventral nucleus of the ventral telencephalon (Vv) a homolog of the LS. In the midbrain, the lateral tuberal nucleus (VTn/NLT) a homolog of the AH, and the anterior tuberal nucleus (ATn) a homolog of the VMH. Figure 1 illustrates the different brain regions analyzed and their relative location along the brain.

**Figure 1:**
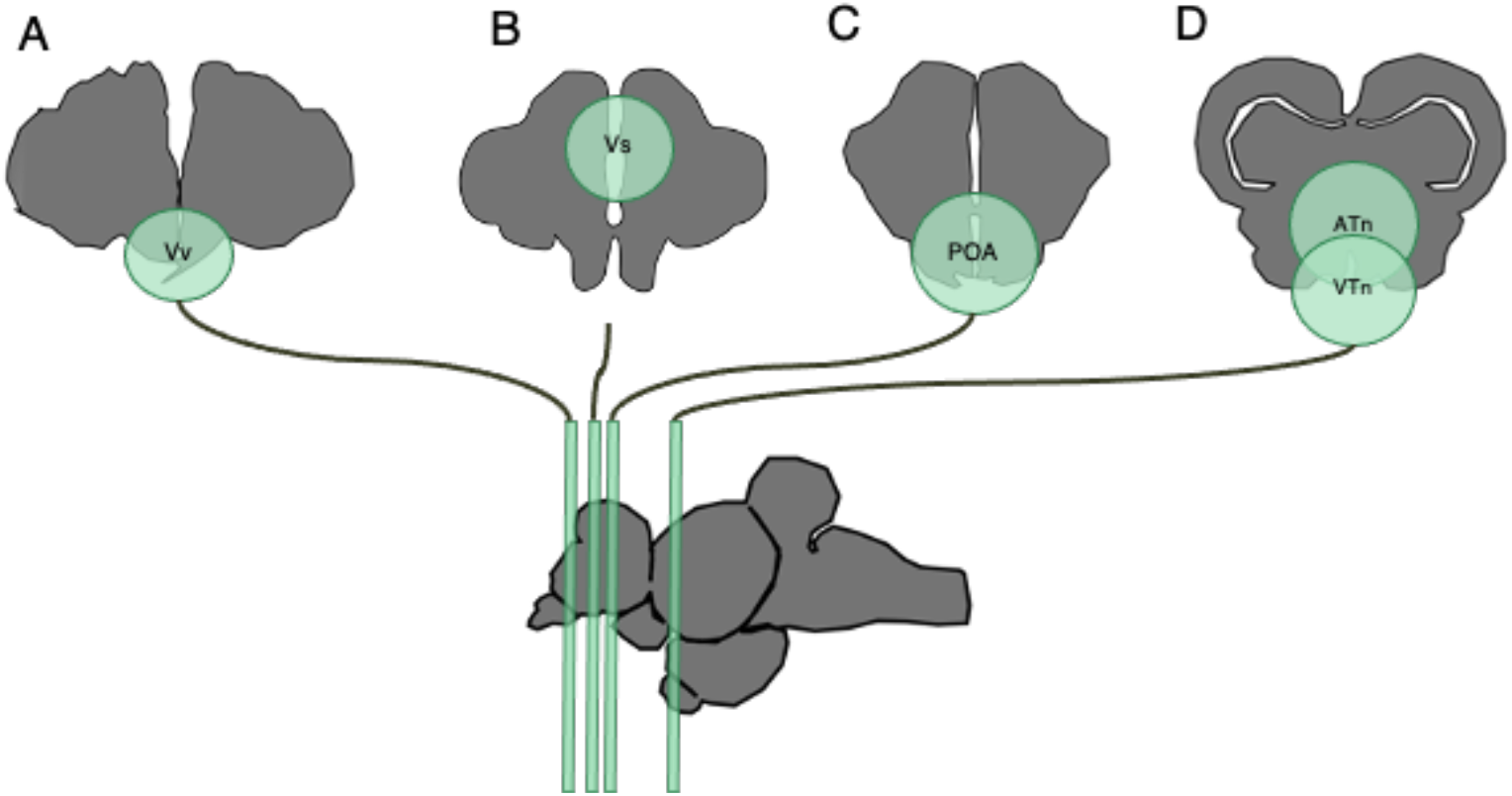
*A. burtoni* coronal brain regions analyzed for *in situ* Hybridization and HCR. (A) shows the Vv (homolog of the LS), (B) shows the VS (partial homolog of the BNST), (C) depicts the preoptic area (POA), (D) shows the ATn (VMH homolog), and the VTn (AH homolog).

For certain brain regions across genotypes, tissue folding and loss during the ISH meant some fish could not be included in the analysis of that brain region. For the Vv, only two WT fish could have expression quantified reliably, so this region was excluded from further analysis. For Het and KO brains the number of fish that could be used for quantification of expression in certain brain regions was as low as 2 or 3 for either group. Given that both AR^Het^ and AR^KO^ males do not differ on the behavioral phenotypes relevant to the proposed link to aromatase, they were combined into a single group called AR mutants, abbreviated as AR^Mut^. Unpaired t-tests between these two genotypes were conducted to confirm they did not differ statistically in expression for any brain region quantified (see results). Final sample sizes for analysis of *cyp19a1* expression across brain regions are as follows: Vs: AR^WT^=4, AR^Mut^= 6; POA: AR^WT^= 4, AR^Mut^= 7; VTN: AR^WT^= 4, AR^Mut^= 8; ATN: AR^WT^= 4, AR^Mut^= 9.

Analysis of brain regions was done with the free source program Image J (imagej.nih.gov/ij/; RRID: SCR_003070) which allowed for tabulation of expression in regions of interest throughout the brain. The images were first converted to 8-bit black and white images and a region of interest (ROI) was drawn surrounding the specific brain region of interest. Once an ROI was selected, the background of the 8-bit images was removed by changing the image threshold using the auto-threshold function on Image J and consistently choosing the same threshold settings for mutant or WT brains. Subsequently, the resulting image was subtracted from the original 8-bit picture resulting in a .tiff image file with prominent staining and low/ non-present background (hereby called the optimal image). The ROI was once again opened, and the threshold of the optimal image was once again adjusted using the threshold function on ImageJ. The thresholded particles within the ROI of the optimal image were then analyzed and numerical values were obtained detailing the intensity and area of each of the particles within the ROI using the analyze particles function on ImageJ. These values were then summed for each hemisphere of the ROI giving the resulting numerical values for each brain region.

### Statistics

Statistical analyses were performed using PRISM software (GraphPad Prism version 9.0) to compare *cyp19a1* expression in mutant and wild type brains. Specifically, unpaired two-tailed t-tests were performed to compare *cyp19a1* signal intensity and area of intensity. If equality of variance or normality assumptions for parametric statistics were not met, log transforms were used to meet those assumptions. Differences were considered significant at p≤0.05.

## Results

### ARα regulates *cyp19a1* expression in the VMH of male *A. burtoni*

Wild-type males, however, showed significantly higher *cyp19a1* expression intensity compared to mutants in the ATn (t (11) =2.538, p=0.0309; Figure 2). There were no significant differences in *cyp19a1* expression of in the other brain regions analyzed. For instance, the expression intensity in Vs (t (8) =0.1154, p=0.9110), the POA (t (9) =0.05026, p=0.6910), and VTn (t (11) =0.08883,p=0.9303; Figure S1) did not differ between wild-type and mutants. No significant differences between wild-type and mutants in the area of intensity of *cyp19a1* expression were detected in all brain regions analyzed (Supplementary file, Figure S2).

**Figure 2:**
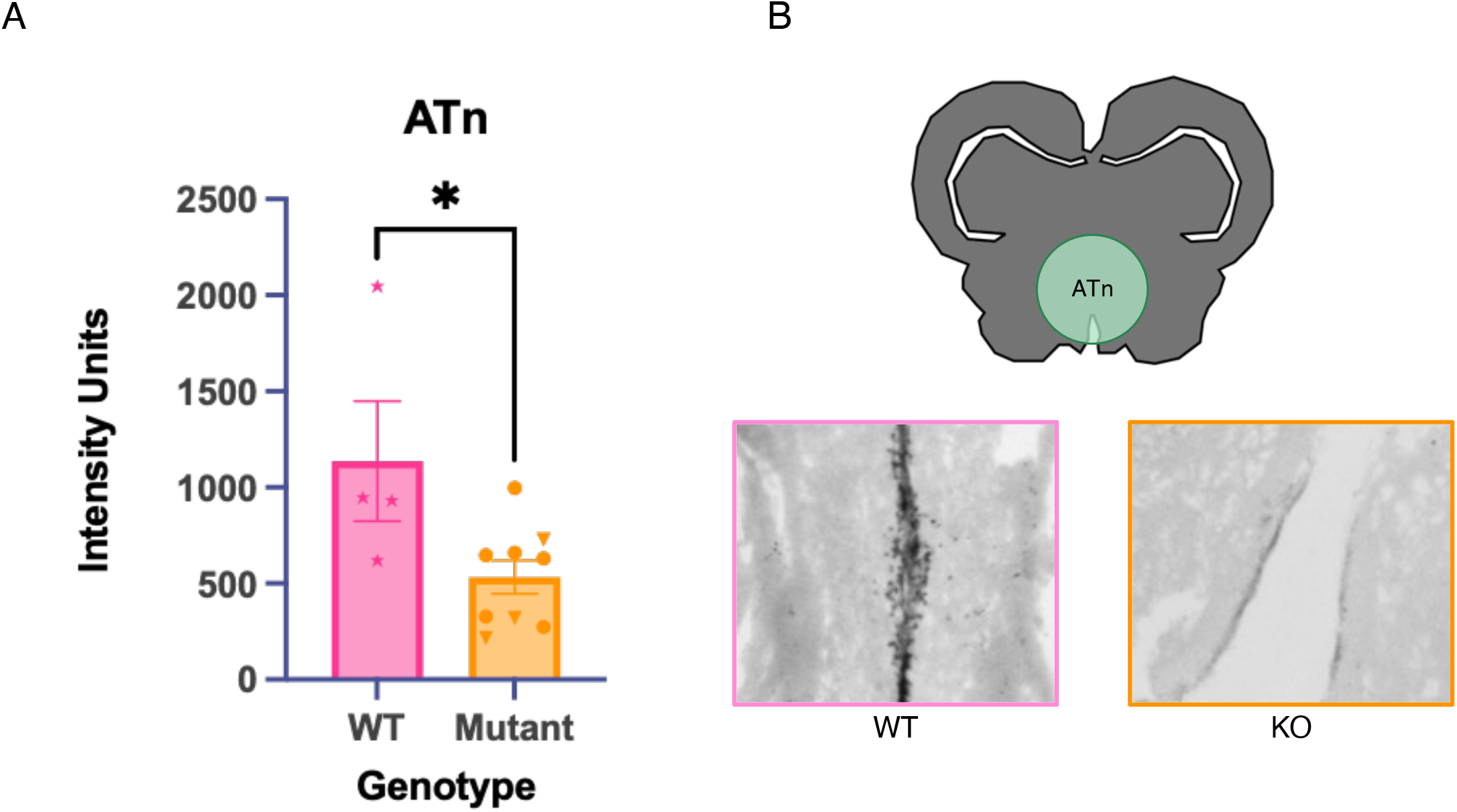
(A) Shows the difference between WT and mutant *cyp19a1* expression (graphed as intensity) in the ATn (VMH). Bars represent mean ± Standard error of the mean (SEM). Stars in the WT group represent each individual fish analyzed. The mutant group represents AR^het^ fish as circles, and AR^KO^ individuals as triangles. (B) Shows a representative image of WT vs mutant brain expression of *cyp19a1* using *in situ* hybridization in the ATn.

### Control of *cyp19a1* expression by ARα is likely indirect

HCR visualization of *cyp191a* expression along with ARα or ARβ expression showed that while ARα and ARβ are both expressed in close proximity of *cyp19a1*^+^ cells, co-expression of either AR with *cyp19a1* is extremely rare even when viewed with z-stacks. Indeed, only in very few instances is either AR found within the same cells. This is true for both the male and female brains analyzed (See Figure 3 for male ATn; See Supplementary file Figures S3-S4 for these patterns in other relevant male regions and in female brains). In addition, similar to previous work *cyp19a1* expression is restricted to the ventricular areas of the brain in both males and females with very little expression in areas other than the ventricles^32^. Qualitatively, *cyp191a* expression seemed to differ between males and females, with females showing higher levels of *cyp191a* expression compared to males, but we did not pursue this potential difference further because it is out of scope of the current study.

**Figure 3:**
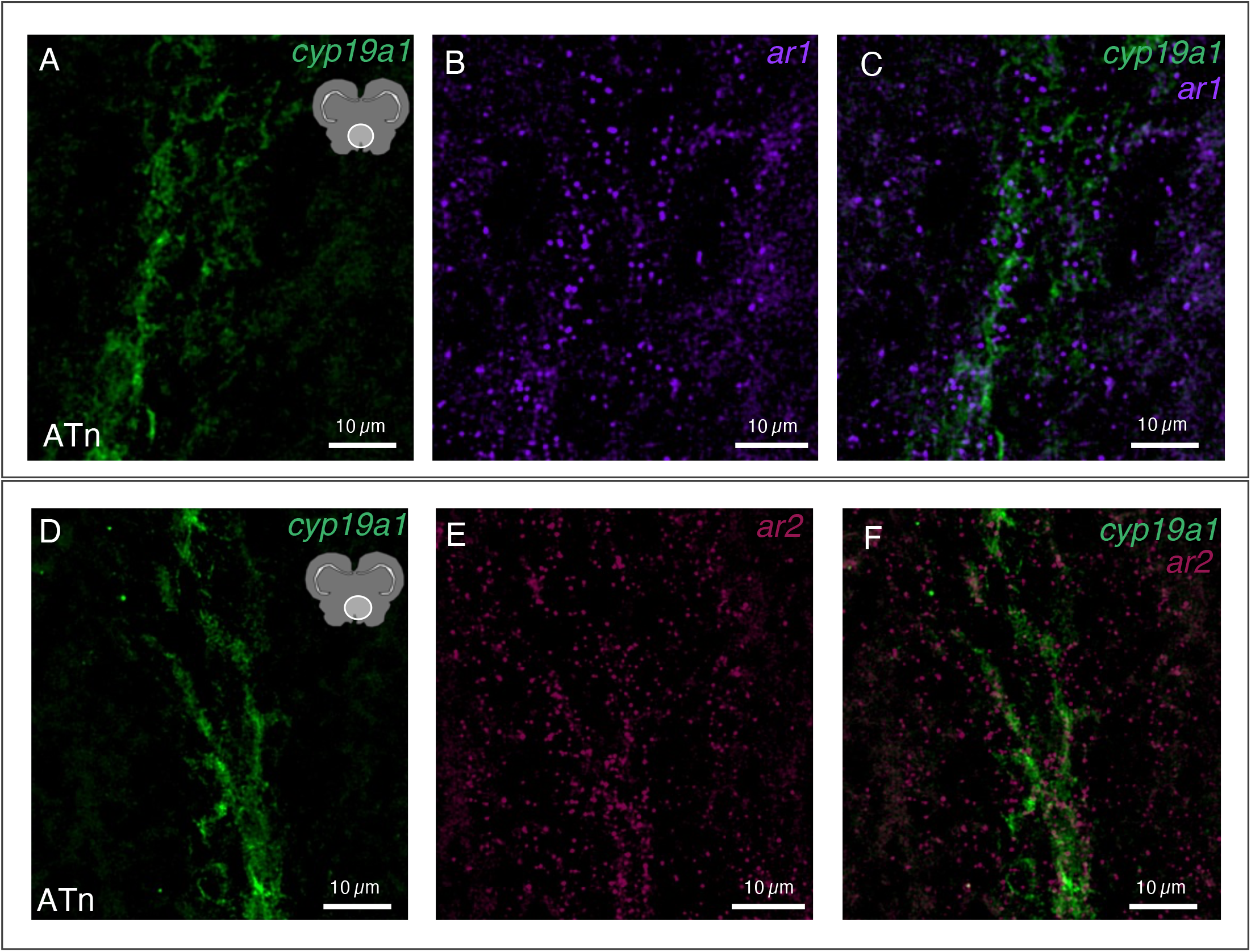
(A-C): Show *cyp19a1* and *ar1* expression in the male ATn. Individual purple puncta represent *ar1* mRNA, while green clusters along the ventricular regions of the brain represent *cyp19a1* mRNA. Note that there is no co-expression of both *ar1* and *cyp19a1* within the same cells. (D-E) Show *cyp19a1* and *ar2* expression. Once again, no co-expression is observed, with individual magenta puncta denoting *ar2* mRNA and *cyp19a1* expression in green along the ventricle.

Expression of ARα and ARβ followed similar patterns compared to previous work^31^. ARα appeared to be mostly concentrated in the anterior part of the brain including the Vv, Vs, POA, ATn, and VTn (See supplementary file Figures S3-S4). Likewise, ARβ shows higher expression patterns around the anterior part of the brain before reducing around the VTn (Figure S3 A-X), and once again showed up in the posterior parts of the brain (not pictured). Furthermore, ARα and ARβ signal appeared to be more sparse than *cyp191a1*, whose expression appeared much more densely.

## Discussion

In the current study we sought to determine if ARα modulates the expression of *cyp19a1* in the male *A. burtoni* brain. Given that ARα and estrogens regulate aggression in *A. burtoni*^14–16^ and androgen signaling enhances aromatase expression in other species^20,21^, we hypothesized that male *A. burtoni* with mutant ARα would possess lower *cyp19a1* expression in regions of the SBN. We found that males deficient in ARα have lower *cyp19a1* expression specifically in the ATn, a homolog of the mammalian VMH. These differences in *cyp19a1* expression are particularly interesting since the VMH is involved in the production of aggressive behavior across different species. Given the conserved functions of both aromatase and the VMH in controlling aggression^18,33–38^, we speculate that ARα governs aggressive displays in male *A. burtoni* through the modulation of *cyp19a1* expression in the ATn ^7,8^.

Extensive research suggests the VMH is critical for the production of aggressive and reproductive behaviors^37–40^. In zebrafish and *A. burtoni*, it has been shown that ARs modulate aggression ^14,41^. The present findings in *A. burtoni* lacking ARα are thus particularly intriguing considering the important role of the VMH and ARs in the production of behavior in other vertebrate species. Yet, available findings have not tested the role of AR signaling in the VMH in the control of male-typical aggression or aromatase expression in teleost or non-teleost vertebrate species. Instead, extensive evidence indicates estrogenic signaling within the VMH is critical for the production of aggression in both male and female mice ^37,38^. Dugger et al^42^ found that AR mutant male mice, which perform significantly fewer aggressive and mating acts compared to WT males, possessed a feminized VMH as measured by volume of this brain region. As mentioned above, androgens increase aromatase expression in male song sparrows^21^ and male rats^22^. Combined with our findings, it is reasonable to speculate that androgen signaling may serve a conserved function across vertebrate taxa to enhance aromatase expression in the VMH, which in turn drives male-typical aggression. This hypothesis is testable and warrants further investigation.

Additionally, our co-expression results are consistent with previous findings from Forlano et. al^43^ which show that AR mRNA and aromatase mRNA/protein are produced in the same brain regions, but not always the same cells, throughout the brain of the midshipman fish (*Porichthys notatus*).

Collectively, this suggests that ARα does not act directly within *cyp19a1*^+^ cells to increase *cyp19a1* expression in *A. burtoni*. Instead, given the consistent close proximity of ARα^+^ cells to *cyp19a1*^*+*^ cells, ARα may increase *cyp19a1* expression through an indirect mechanism, such as a transsynaptic or transcellular process^44,45^. As *cyp19a1* expression is assumed to be nearly solely localized to glia cells in *A. burtoni*^32^, this hypothesis would assume a neuron-glia interaction, or potentially a glia-glia interaction. Either interaction is hypothetically possible since it is not known which cell-types in the brain ARα is present in *A. burtoni*, but in other species AR is present in both neurons and glial cells^46–48^. Another mechanism could be developmentally dependent. For example, during earlier stages of development, it may be that ARα acts within the same cells in which it enhances *cyp19a1* expression, an effect that endures until adulthood, a time during which ARα is no longer expressed in those cells. Both ARs and *cyp19a1* are expressed in zebrafish as early as 24-48 hours post fertilization for *cyp19a1* ^49^ and 3-5 days for AR^50^ in zebrafish, meaning it is plausible AR could influence *cyp19a1* expression in the developing teleost brain. These hypotheses lay the groundwork for future developmental, anatomical, and functional studies in *A. burtoni* and other teleosts to further dissect how androgenic and estrogenic signaling shape social behaviors.

## Conclusions

Overall, our findings showed that ARα deficiency reduces *cyp19a1* expression in the *A. burtoni* VMH. This is an intriguing finding when considering the importance of the VMH in the production of aggressive behavior across different vertebrate species and suggests that appropriate levels of *cyp19a1* may be an important component in the production aggressive and reproductive behaviors in males of our species. However, many unknowns remain including the specific ways in which ARα indirectly influences *cyp19a1* expression. Future studies should also aim to investigate whether the second AR paralog in *A. burtoni* (ARβ) has a different effect in the expression of *cyp19a1* in regions of the SBN. Furthermore, *cyp19a1* expression patterns should be analyzed in females lacking ARα to further elucidate the fundamental role of androgens in the production of social behavior in *A. burtoni*. Future studies on novel teleost steroid signaling genes will enrich our understanding of the molecular and neural mechanisms that contribute to the generation of behavior.

## Acknowledgements

We thank Melanie Dussenne and Kathleen Munley for valuable suggestions on the HCR figures and Caitlin Kennedy for helpful discussions on the ISH procedures.

