## Supplementary Material for "Androgen receptor alpha regulates aromatase expression in the ventromedial hypothalamus of male cichlids"

**Author’s contributions:** B.A.A. and M.S.L. designed the study; M.S.L. performed the experiments, collected the data, analyzed the data, and made the figures. B.A.A. and M.S.L. wrote and edited the manuscript.

**Funding:** This research was supported by a Beckman Young Investigator Award from the Arnold and Mabel Beckman Foundation, an NIH grant R35GM142799, and a University of Houston-National Research University Fund startup R0503962 to B.A.A.

**Acknowledgements:** We thank Melanie Dussenne and Kathleen Munley for valuable suggestions on the HCR figures and Caitlin Kennedy for helpful discussions on the ISH procedures.

**Competing interest**: The authors declare no competing or financial interests.

### Abstract

Social behaviors are regulated by sex steroid hormones such as androgens and estrogens. However, the specific molecular and neural processes modulated by steroid hormones to generate social behaviors remain to be elucidated. We investigated whether some actions of androgen signaling in the control of social behavior may occur through the regulation of estradiol synthesis in the highly social cichlid fish *Astatotilapia burtoni*. Specifically, we examined the expression of *cyp19a1,* a brain-specific aromatase, in the brains of male *A. burtoni* lacking a functional AR$\alpha$ gene, which was recently found to be necessary for aggression in this species. We found that *cyp19a1* expression is higher in wild-type males compared to AR$\alpha$ mutant males in the homolog of the mammalian ventromedial hypothalamus (VMH), a brain region that governs aggression across taxa. Using *in situ* hybridization chain reaction (HCR), we determined that AR$\alpha^+$ and *cyp19a1*$^+$ cells are commonly nearby, but very infrequently was co-expression observed, including in the fish VMH. We speculate that AR$\alpha$ may modulate *cyp19a1* expression in the fish VMH to govern aggression through an indirect, potentially transsynaptic, mechanism. These studies provide novel insights into the hormonal mechanisms of social behavior and lay a foundation for future functional studies.

### Supplementary Results

#### ***AR$\alpha$ regulates cyp19a1 expression in the VMH of male A. burtoni***

The area of detectable expression for *cyp19a1* was also graphed for each of the WT vs. Mutant groups. For this measure, no significant results were found for any of the brain regions (Supplementary Figure 2). The Vs showed no difference between the WT and mutant areas of *cyp19a1* expression (t (8) =0.07124, p=0.9450). Similarly, the POA showed no differences in expression between the WT and mutant groups (t (9) =1.674), p= 0.1284). The ATn and VTn respectively both showed no differences between the area of expression of the WT and mutant groups (t (11) =1.376, p=0.1960 and t(11)= 0.6513, p=0.5283 respectively; Supplementary Figure 2).

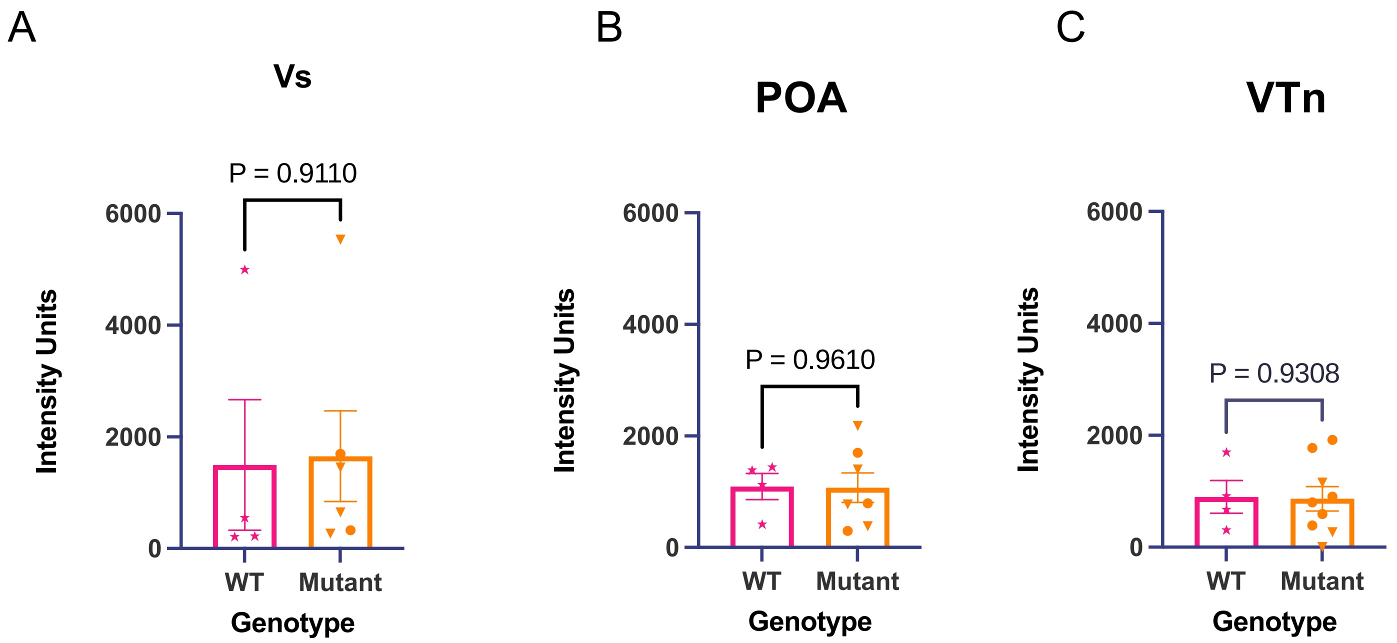


*Figure 1:* (A) shows the difference between WT and mutant *cyp19a1* expression (graphed Intensity *of expression)* in the *Vs (LS homolog)*. (B) Shows the differences between WT and mutant *cyp19a1 group expression in the POA (preoptic area). (C) Depicts intensity differences between the WT and mutant fish in the* VTn (AH homolog). Bars represent mean ± Standard error of the mean (SEM). Stars in the WT group represent each individual fish brains analyzed. The mutant group represents AR $^{het}$ fish as circles, and AR$^{KO}$ individuals as triangles.

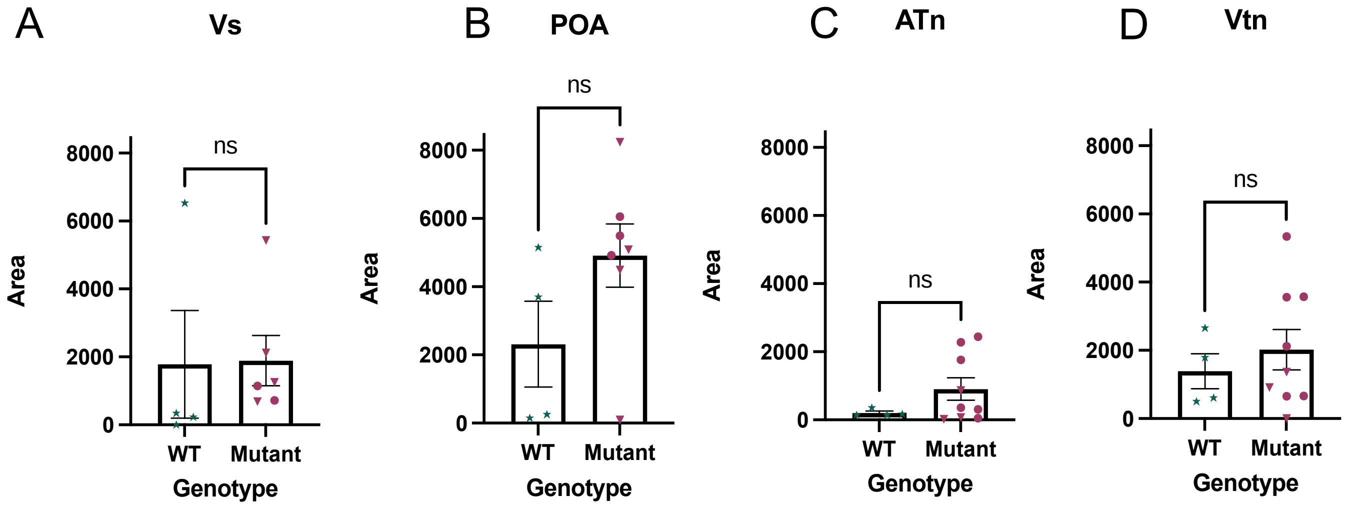


*Figure 2:* (A) shows the difference between WT and mutant *cyp19a1* area of expression in the *Vs (LS homolog)*. (B) Shows the differences between WT and mutant *cyp19a1 group area of expression in the POA (preoptic area). (C) Depicts intensity differences between the WT area and mutant fish area in the ATn* (VMH homolog). *(D) Depicts area differences between the WT and mutant fish in the* VTn (AH homolog). Bars represent mean ± Standard error of the mean (SEM). Stars in the WT group represent each individual fish brains analyzed. The mutant group represents AR $^{het}$ fish as circles, and AR$^{KO}$ individuals as triangles.

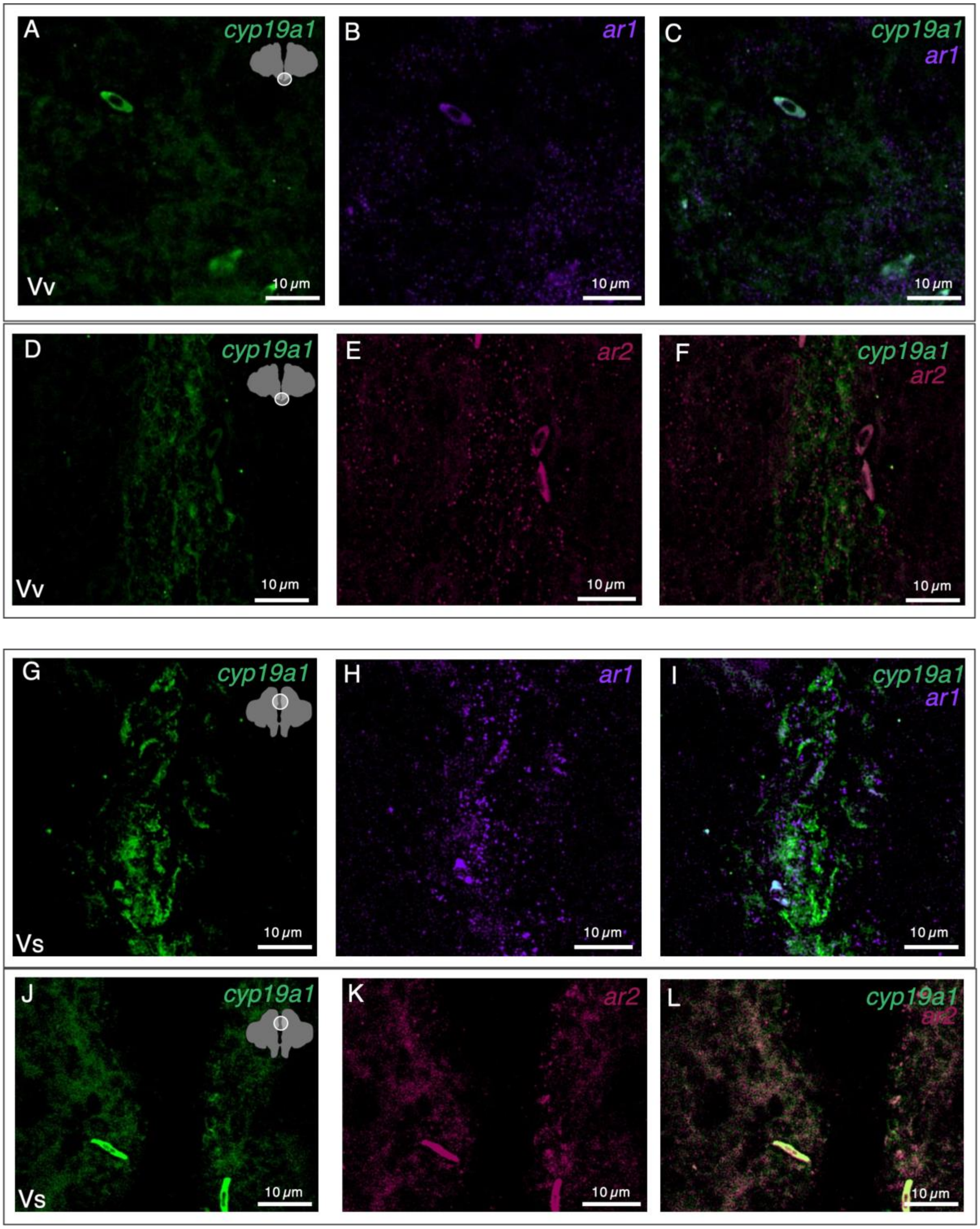
A
cyp19a1
Vv
10 µm
B
ar1
10 µm
C
cyp19a1
ar1
10 µm
D
cyp19a1
Vv
10 µm
E
ar2
10 µm
F
cyp19a1
ar2
10 µm
G
cyp19a1
Vs
10 µm
H
ar1
10 µm
I
cyp19a1
ar1
10 µm
J
cyp19a1
Vs
10 µm
K
ar2
10 µm
L
cyp19a1
ar2
10 µm

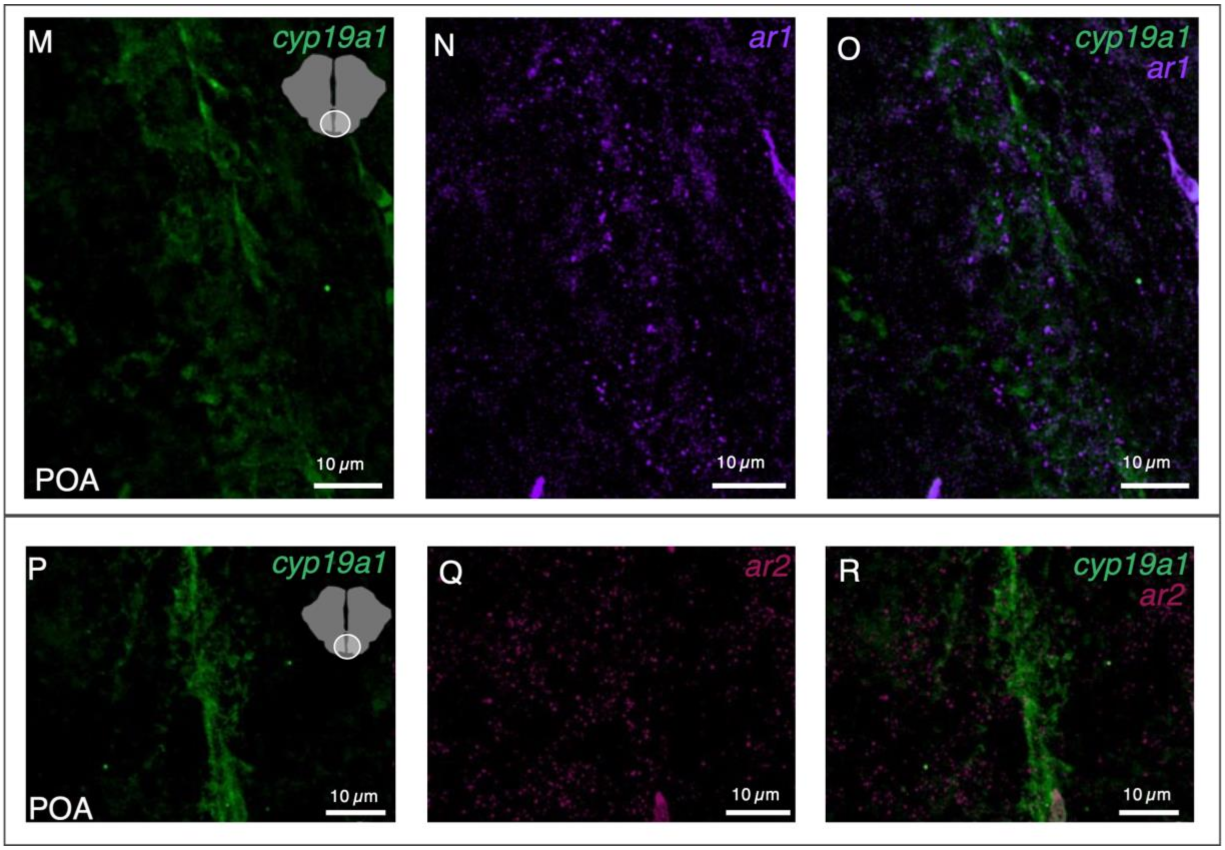
M
cyp19a1
POA
10 μm
N
ar1
10 μm
O
cyp19a1
ar1
10 μm
P
cyp19a1
POA
10 μm
Q
ar2
10 μm
R
cyp19a1
ar2
10 μm

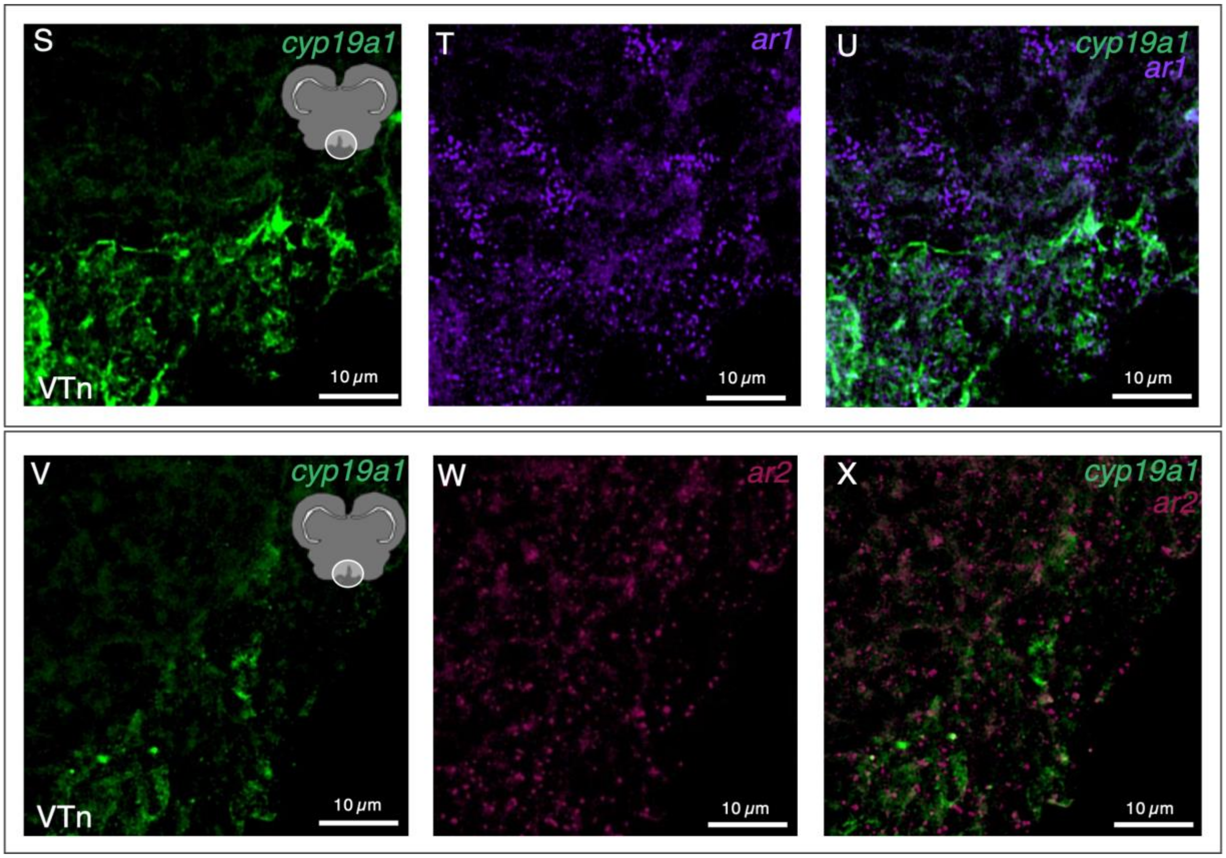


*Figure 3:* (A-C) Show VV. Individual purple puncta represent *ar1* mRNA, while green clusters along the ventricular regions of the brain represent *cyp19a1* mRNA. Note that there is no co-expression of both *ar1* or *cyp19a1* within the same cells and *cyp19a1* expression seems lower than subsequent panels. (D-F) Show *cyp19a1* and *ar2* expression. Once again, no co-expression is observed, with individual magenta puncta denoting *ar2* mRNA and cyp*19a1* expression in green along the ventricle.
(G-I) Show *cyp19a1* and *ar1* expression in the male Vs with most *cyp19a1* expression concentrated along the ventricles with nearby puncta of *ar1* expression. Note there is no co-expression of both genes within this region. (J-L)Once again depict the male Vs with *cyp19a1* (in green) and *ar2* (magenta) expression. Note both genes are very sparsely spread along this region and some areas of co-expression may exist but background signal from both genes makes co-expression difficult to interpret. (M-O) Illustrate the male POA with *cyp19a1* and *ar1* expression. Note the lack of co-expression of the two genes. (P-R) Similarly show the POA region with *ar2* and *cyp19a1* here it is notable that *ar2* expression seems to be fainter and sparser compared to panels M-O in the same region. (S-U) Depict VTn region of the male brain with *cyp19a1* and *ar1* expression. Note that will expression of both *ar1* and *cyp19a1* are quite dense in this region compared to others in previous panels. (V-X) Show *ar2* and *cyp19a1* expression in VTn with lower signal expression from *ar2* and *cyp19a1* compared to panels S-U. Note that for both *ar1* and *ar2* there is no co-expression observed in *cyp19a1* positive cells within the VTn.

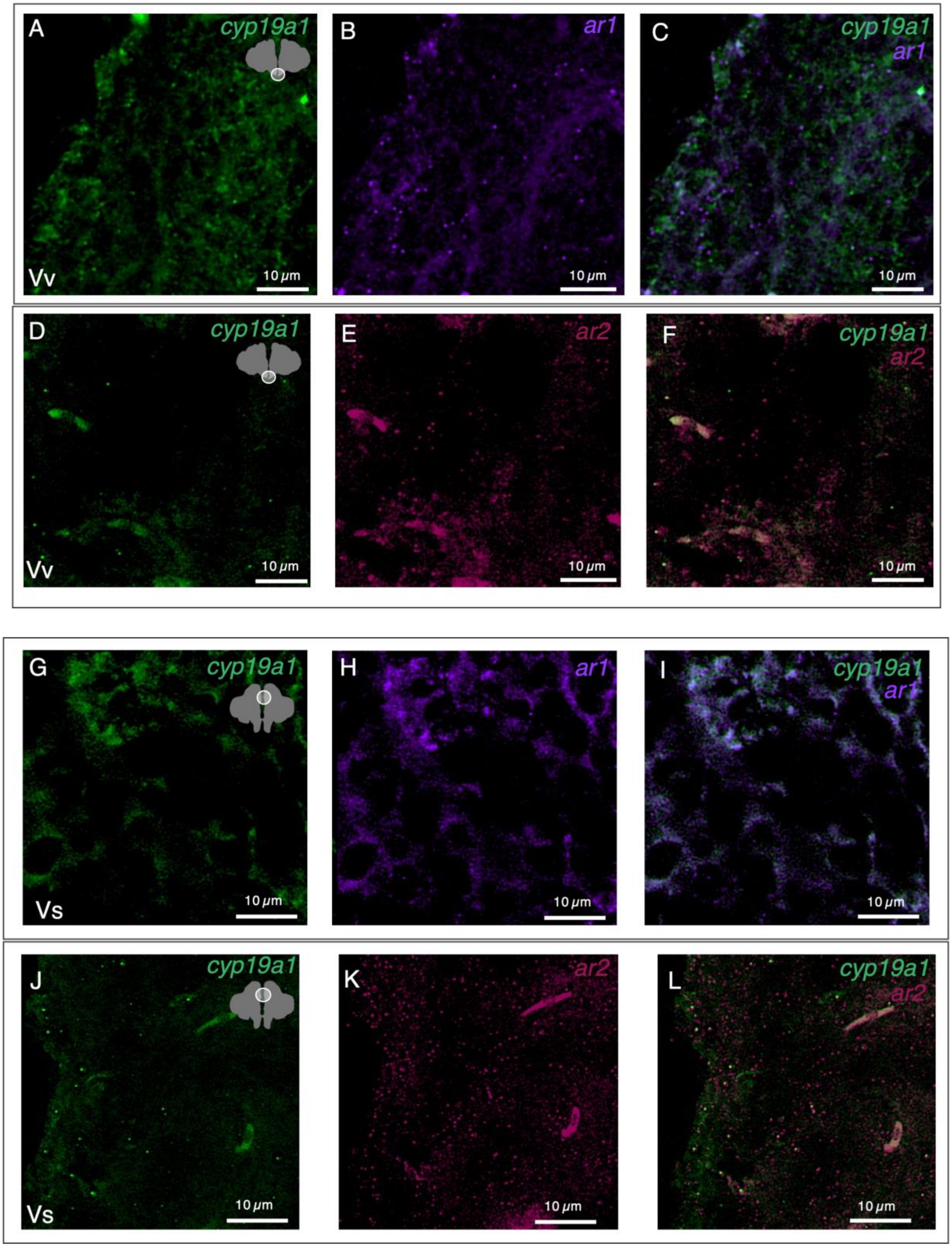
A
cyp19a1
Vv
10 μm
B
ar1
10 μm
C
cyp19a1
ar1
10 μm
D
cyp19a1
Vv
10 μm
E
ar2
10 μm
F
cyp19a1
ar2
10 μm
G
cyp19a1
Vs
10 μm
H
ar1
10 μm
I
cyp19a1
ar1
10 μm
J
cyp19a1
Vs
10 μm
K
ar2
10 μm
L
cyp19a1
ar2
10 μm

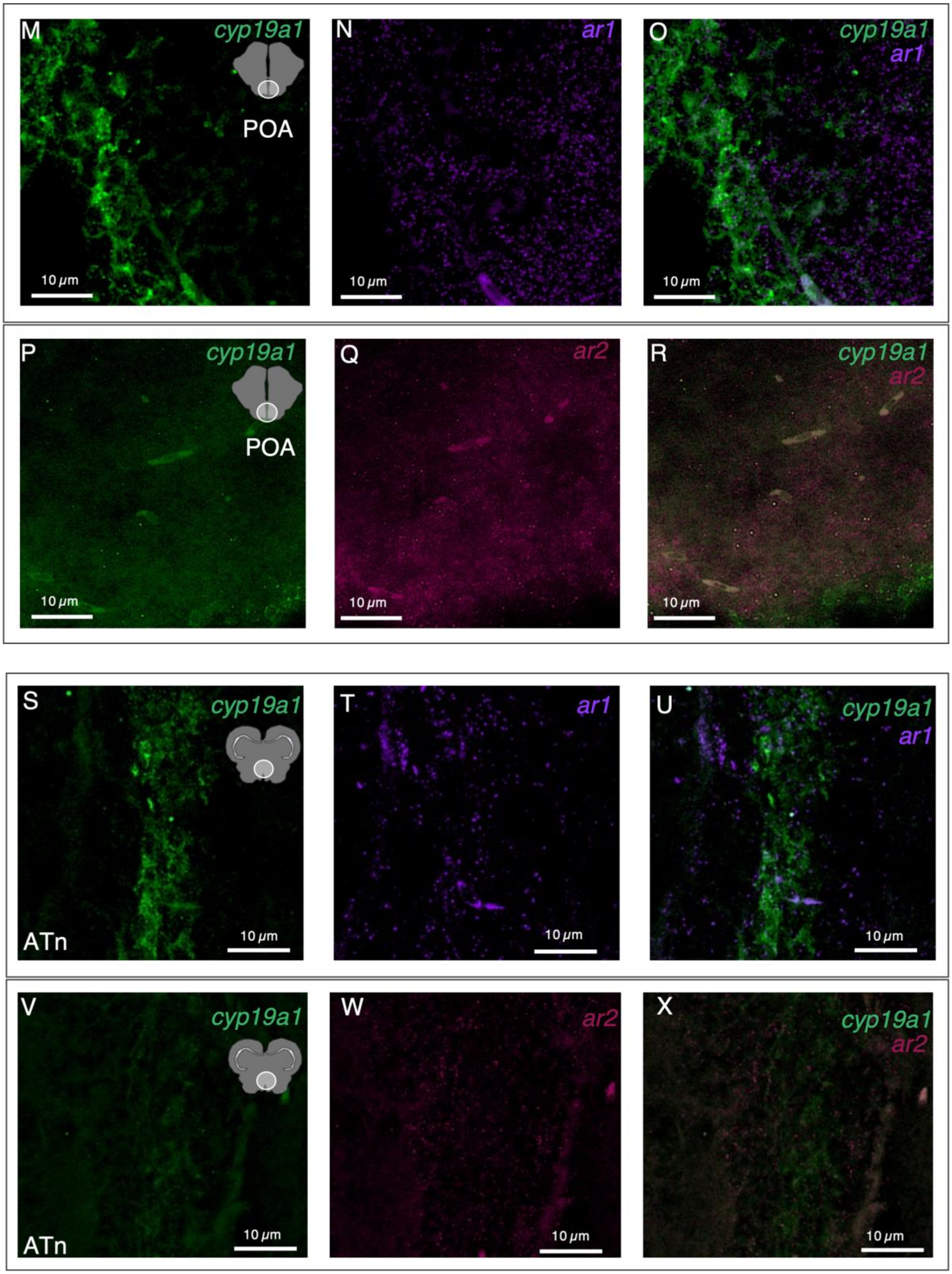
M
cyp19a1
POA
10 μm
N
ar1
10 μm
O
cyp19a1
ar1
10 μm
P
cyp19a1
POA
10 μm
Q
ar2
10 μm
R
cyp19a1
ar2
10 μm
S
cyp19a1
ATn
10 μm
T
ar1
10 μm
U
cyp19a1
ar1
10 μm
V
cyp19a1
ATn
10 μm
W
ar2
10 μm
X
cyp19a1
ar2
10 μm

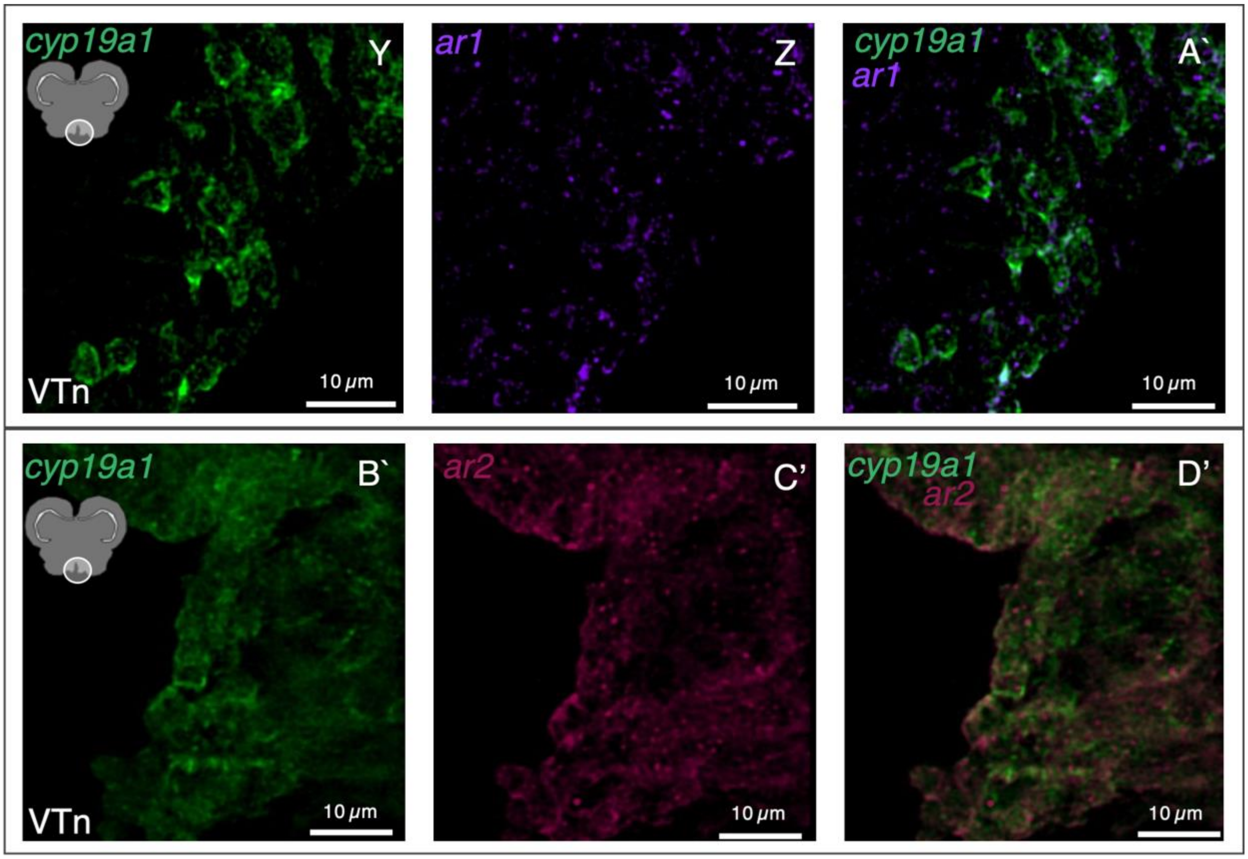


*Figure 4:* (A-C) Show the female VV. Individual purple puncta represent *ar1* mRNA, while green clusters along the ventricular regions of the brain represent *cyp19a1* mRNA. Note that there is no co-expression of both *ar1* and *cyp19a1* within the same cells. (D-F) Show *cyp19a1* and *ar2* expression. Once again, no co-expression is observed, with individual magenta puncta denoting *ar2* mRNA and cyp*19a1* expression in green along the ventricle. Further expression of both *ar2* seems sparse within this region.
(G-I) Show *cyp19a1* and *ar1* expression in the female Vs with note that expression of both *cyp19a1* and *ar1* seem sparse and thus co-expression is hard to gauge from signal background. (J-L) Once again depict the male Vs with *cyp19a1* (in green) and *ar2* (magenta) expression. Note both genes are very sparsely spread along this region, but *ar2* seems to have slightly higher expression compared to *ar1* in J-L. (M-O) Illustrate the female POA with *cyp19a1* and *ar1* expression. Note the lack of co-expression of the two genes, but apparent proximity of both. (P-R) Similarly show the POA region with *ar2* and *cyp19a1,* here it is notable that *ar2* expression seems to be fainter and sparser compared to panels M-O in the same region. (S-U) Depict ATn region of the female brain with *cyp19a1* and *ar1* expression. Note that will expression of both *ar1* and *cyp19a1* are quite sparse in this region compared to previous panels. (V-X) Show *ar2* and *cyp19a1* expression in ATn with lower signal expression from *ar2* and *cyp19a1* compared to panels S-U. Note that for both *ar1* and *ar2* there is no co-expression observed in *cyp19a1* positive cells within the ATn. (Y-A') Show the VTn of a female with *ar1* and *cyp19a1*. Note here the distinct morphology of the *cyp19a1* positive cells and how they compare to both previous panels and panels B'-D'. Once again there is no observable co-expression of both genes. (B'-D') Show the same VTn region with *ar2* and *cyp19a1*, note that both genes seemed to be expressed in the same regions and perhaps in some of the same cells but it is still very difficult to tell.
